# Chromatin structure-dependent histone incorporation revealed by a genome-wide deposition assay

**DOI:** 10.1101/641381

**Authors:** Hiroaki Tachiwana, Mariko Dacher, Kazumitsu Maehara, Akihito Harada, Yasuyuki Ohkawa, Hiroshi Kimura, Hitoshi Kurumizaka, Noriko Saitoh

**Affiliations:** Division of Cancer Biology, The Cancer Institute of Japanese Foundation for Cancer Research, Tokyo 135-8550, Japan; Laboratory of Chromatin Structure and Function, Institute for Quantitative Biosciences, The University of Tokyo, Tokyo 113-0032, Japan; Division of Transcriptomics, Medical Institute of Bioregulation, Kyushu University, Fukuoka 812-0054, Japan; Cell Biology Center, Institute of Innovative Research, Tokyo Institute of Technology, Yokohama 226-8503, Japan

## Abstract

In eukaryotes, histone variant distribution within the genome is the key epigenetic feature. To understand how each histone variant is targeted to the genome, we developed a new method, in which epitope-tagged histone complexes are introduced into permeabilized cells and incorporated into their chromatin. We found that the incorporation of histones H2A and H2A.Z mainly occurred at less condensed chromatin (open), suggesting that the condensed chromatin (closed) is a barrier for histone incorporation. To overcome this barrier, H2A, but not H2A.Z, uses a replication-coupled deposition mechanism. This led to the recapitulation of the pre-existing chromatin structure: the genome-wide even distribution of H2A and the exclusion of H2A.Z from the closed chromatin. Intriguingly, an H2A.Z mutant with mutations in the developmentally essential region was incorporated into closed chromatin. Our study revealed that the combination of chromatin structure and DNA replication dictates the differential histone deposition for maintaining the epigenetic chromatin states.

## Introduction

Eukaryotic genomic DNA is packaged into chromatin, in which the basic structural unit is the nucleosome. The nucleosome is composed of around 150 base pairs of DNA and a histone octamer consisting of two copies of each core histone, H2A, H2B, H3, and H4 (Luger, Mäder, Richmond, Sargent, & Richmond, 1997). Chromatin is not only the storage form of genomic DNA but also the regulator of DNA-templated processes, such as transcription, replication, repair, and chromosome segregation. To enable these processes, chromatin forms various structures that can be reversibly altered. Historically, cytological studies first identified euchromatin and heterochromatin, which are transcriptionally active and inactive regions, respectively (Passarge, 1979). Subsequent studies revealed that euchromatin coincides with nuclease hypersensitivity, indicating that it forms a more accessible structure (open chromatin) than heterochromatin (closed chromatin) (Garel & Axel, 1976; Spiker, Murray, & Thompson, 1983; Tsompana & Buck, 2014; Weintraub & Groudine, 1976). Although these chromatin structures are involved in the regulation of DNA template-mediated processes, little is known about how the open and closed chromatin configurations are formed and maintained.

Among the chromatin associated proteins, histones have a significant impact on the chromatin structure. In humans, the canonical histones, H2A, H2B, and H3.1, have non-allelic variants with distinct expression and/or localization patterns (Buschbeck & Hake, 2017; Maehara et al., 2015). Canonical H3.1 and H2A are expressed in S phase and show genome-wide localizations (Buschbeck & Hake, 2017; Wu, Tsai, & Bonner, 1982). In contrast, the histone variants H3.3 and H2A.Z are expressed throughout the cell cycle and concentrated at promoters in both open chromatin and pericentric heterochromatin (Ahmad & Henikoff, 2002; Boyarchuk, Filipescu, Vassias, Cantaloube, & Almouzni, 2014; Drane, Ouararhni, Depaux, Shuaib, & Hamiche, 2010; Goldberg et al., 2010; Greaves, Rangasamy, Ridgway, & Tremethick, 2007; Jin & Felsenfeld, 2007; Raisner et al., 2005; Rangasamy, Berven, Ridgway, & Tremethick, 2003; Sarcinella, Zuzarte, Lau, Draker, & Cheung, 2007; Wu et al., 1982). In addition, H3.3 localizes at telomeres (Goldberg et al., 2010). Another H2A variant, MacroH2A, is localized at transcriptionally suppressed chromatin, such as inactive X chromosomes (Costanzi, Stein, Worrad, Schultz, & Pehrson, 2000). Along with the canonical H2A, the localization of H2A.X, which functions in DNA repair, is genome-wide (Yukawa et al., 2014). Thus, in spite of the high sequence homology between the canonical and variant forms, each histone has specific functions. A previous study identified the six essential residues responsible for the H2A.Z-specific functions, which are located in the αC helix called the M6 region (Clarkson, Wells, Gibson, Saint, & Tremethick, 1999). The H2A.Z swap mutant, in which M6 was replaced with the equivalent residues in H2A, failed to rescue the embryonic lethality of the H2A.Z null mutant in *Drosophila melanogaster*. Thus, the importance of M6 in the organism was evident; however, its molecular mechanism remained elusive.

Histone deposition and exchange have been studied for more than 30 years (Clément et al., 2018; Jackson, 1990; Jansen, Black, Foltz, & Cleveland, 2007; Kimura & Cook, 2001; Louters & Chalkley, 1985; Ray-Gallet et al., 2011; Tachiwana et al., 2010). An *in vivo* histone deposition study must distinguish the pre-incorporated histones (parental histones) from the newly incorporated histones (new histones). To enable this, initial studies used radio-labeled histones and revealed that histone deposition occurred by both replication-coupled and replication-independent mechanisms (Jackson, 1990; Louters & Chalkley, 1985). Several key proteins involved in these mechanisms have been identified. Anti-silencing function 1 (ASF1), together with chromatin assembly factor 1 (CAF-1), plays indispensable roles in replication-coupled H3.1 deposition (Smith & Stillman, 1989; Tagami, Ray-Gallet, Almouzni, & Nakatani, 2004; Tyler et al., 1999; Tyler, Bulger, Kamakaka, Kobayashi, & Kadonaga, 1996; Tyler et al., 2001). CAF-1 directly interacts with proliferating cell nuclear antigen (PCNA), leading to the assembly of H3.1 at replicating chromatin (Moggs et al., 2000; Shibahara & Stillman, 1999). The histone regulator A (HIRA) or death associated protein (DAXX) promotes the accumulation of H3.3 at transcription sites and regulatory elements in a replication-independent manner (Drane et al., 2010; Goldberg et al., 2010; Ray-Gallet et al., 2002; 2011; Tagami et al., 2004). H2A- and H2A.Z-specific chaperones have also been identified (Obri et al., 2014). Facilitates chromatin transcription (FACT), a chromatin remodeling complex, may be involved in replication-coupled H2A deposition (Orphanides, LeRoy, Chang, Luse, & Reinberg, 1998; Ransom, Dennehey, & Tyler, 2010; Wittmeyer & Formosa, 1997). ANP32E functions in H2A.Z eviction from chromatin, and the SNF2-related CBP activator protein (SRCAP) chromatin-remodeling subunit YL1 promotes H2A.Z deposition at promoters (Latrick et al., 2016; Liang et al., 2016; Mao et al., 2014; Obri et al., 2014; Wong, Cox, & Chrivia, 2007).

The precise distribution of histones is crucial for chromatin organization and its epigenetic states. The ChIP-seq analysis method is powerful and useful to visualize steady state histone localizations; however, it does not enable the analysis of the incorporation of each histone into open/closed chromatin or both (genome-wide). To analyze histone incorporations, several approaches using fluorescence imaging, mass spectrometry, inducible tagged proteins, and labeling of newly synthesized proteins have been developed (Deal & Henikoff, 2010). Imaging-based methods, such as fluorescence recovery after photobleaching (FRAP) and SNAP/CLIP-tag technology, were developed to analyze the histone dynamics in living cells. FRAP is suitable for investigating the histone mobility in living cells (Kimura & Cook, 2001; Tachiwana et al., 2010). The SNAP/CLIP-technology is effective for analyzing the histone deposition, as it can distinguish parental histones from new histones in cells with cell-permeable fluorophores that covalently bind to the tag (Clément et al., 2018; Jansen et al., 2007; Ray-Gallet et al., 2011). Although these are powerful methods, they have resolution limitations since they are based on imaging. The SNAP/CLIP-technology led to the establishment of the time-ChIP method, in which a biotin labeled SNAP-histone is captured (Deaton et al., 2016). The time-ChIP method was developed to measure the stability of parental histones, rather than the histone incorporation, as it has a time lag during the synthesis and labeling of histones, which limits the time resolution (Deaton et al., 2016; Siwek, Gómez-Rodríguez, Sobral, Corrêa, & Jansen, 2018). To uncover the correlation between histone incorporation and open/closed chromatin structures, a new analysis method for the histone incorporation at the DNA sequence level is desired.

Permeabilized cells are useful to dissect the molecular pathways in nuclear events (Adam, Marr, & Gerace, 1990; Okuno, Imamoto, & Yoneda, 1993). In assays with such cells, the cellular membranes are permeabilized by a nonionic detergent treatment. In the permeabilized cells, the chromatin and nuclear structures remain intact and react with exogenously added proteins (Kimura et al., 2006; Maison et al., 2002; Misteli & Spector, 1996; Saitoh et al., 2006). A previous study showed that an exogenously added GFP-tagged histone, prepared from cultured human cells, was incorporated into the chromatin of permeabilized cells in the presence of a cellular extract (Kimura et al., 2006). Moreover, permeabilized cells are suitable for monitoring replication timing, by labeling the nascent DNA with exogenously introduced nucleotides (Kimura et al., 2006; Misteli & Spector, 1996).

In the present study, we developed a new method, in which a reconstituted histone complex, instead of a fluorescent protein-tagged histone, was added to permeabilized cells. We named this the RhIP assay (<u>R</u>econstituted <u>h</u>istone complex Incorporation into chromatin of <u>P</u>ermeabilized cell). Since the histone complexes are reconstituted *in vitro* using epitope-tagged recombinant histones, RhIP with sequencing allows the analysis of incorporations at the DNA sequence level, without the need for specific antibodies. We found that the chromatin structure regulates the histone incorporations, which may be necessary for maintaining the epigenetic state of chromatin.

## Results

### RhIP assay reproduces *in vivo* histone deposition

To understand how histones are incorporated into chromatin in cells, we developed the RhIP assay, in which an *in vitro* reconstituted histone complex, nucleotides, and a cellular extract are added to permeabilized cells (Figure 1A). We first confirmed that the RhIP assay can recapitulate the specific histone incorporations observed in cells. The H3.1-H4 incorporation into chromatin is coupled with replication, while the H3.3-H4 incorporation occurs throughout the cell cycle (Ahmad & Henikoff, 2002). We reconstituted H3-H4 complexes *in vitro*, using recombinant H3.1, H3.3, and H4 (Figure 1B). H3.1 and H3.3 were fused to HA and FLAG tags at their C-termini, respectively. The permeabilized cells were then prepared by treating HeLa cells with a nonionic detergent, Triton X-100, and the reconstituted H3-H4 complexes were mixed with the cellular extract and nucleotides. Cy5-dUTP was also added, in order to monitor DNA replication. After the reaction, the exogenously added H3.1 and H3.3 were detected with antibodies against the HA and FLAG tags, respectively (Figure 1C-E). As a result, the H3.1 was detected in the Cy5 positive cells (S phase cells), while the H3.3 was detected irrespective of the Cy5 signal. These results indicate that the reconstituted H3.1-H4 and H3.3-H4 complexes were incorporated into the chromatin of permeabilized cells with the same dynamics observed in cells.

**Figure 1.**
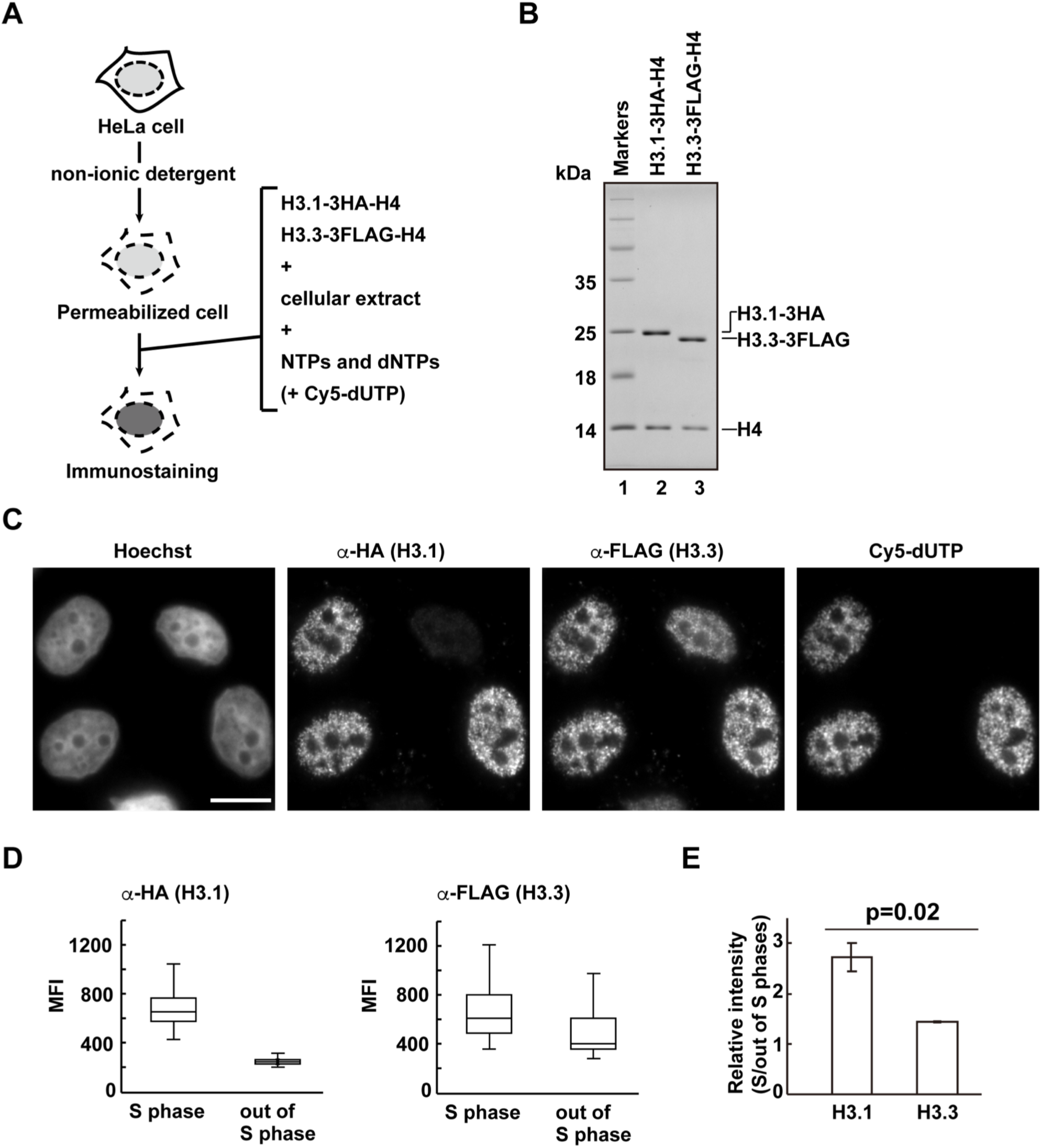
RhIP (<u>R</u>econstituted <u>h</u>istone complex <u>I</u>ncorporation into chromatin of <u>P</u>ermeabilized cells) assay, using H3.1-H4 and H3.3-H4 complexes. (A) Schematic representation of the RhIP assay, using reconstituted H3.1-H4 and H3.3-H4 complexes. Permeabilized cells were prepared from HeLa cells treated with non-ionic detergent, to perforate the cellular membranes. The *in vitro* reconstituted H3-H4 complexes were then added to the cells with the cellular extract and nucleotides. Cy5-dUTP was added to label the nascent DNA, so replication could be monitored. (B) Reconstituted H3.1-H4 and H3.3-H4 complexes were analyzed by SDS-16% PAGE with Coomassie Brilliant Blue staining. The 3HA and 3FLAG tags were fused to the C-termini of H3.1 and H3.3, respectively. Lane 1 indicates the molecular mass markers, and lanes 2 and 3 indicate the H3.1-H4 and H3.3-H4 complexes, respectively. (C) RhIP-immunostaining of H3.1 and H3.3. Exogenously added H3-H4 complexes were stained with an anti-HA or -FLAG antibody. Cells in S phase were monitored with Cy5-dUTP, which was incorporated into the nascent DNA. Bar indicates 10 μm. (D) Quantification of Figure 1C. The mean fluorescence intensities (MFI) of H3.1-3HA (left) and H3.3-3FLAG (right) were measured. Nuclei were divided into S phase (Cy5 positive) and out of S phase (Cy5 negative) (n>150). (E) Relative intensity of H3.1 or H3.3 signal in S phase against signal out of S phase. Experiments were repeated three times and averaged data are shown. The two-tailed Student’s t-test was used for the statistical comparisons.

We next examined whether the cellular extract is essential or replaceable by the histone chaperones, Nap1 or Asf1 (Figure 1-figure supplement 1A). Nap1 and Asf1 bind to the H2A-H2B and H3-H4 complexes *in vivo*, respectively, and both promote nucleosome formation *in vitro* (Ishimi et al., 1984; Munakata, Adachi, Yokoyama, Kuzuhara, & Horikoshi, 2000; Tachiwana, Osakabe, Kimura, & Kurumizaka, 2008; Tyler et al., 1999). Human Nap1 and Asf1 were purified as recombinant proteins (Figure 1-figure supplement 1B). The H3.1-H4 complex was then added to the permeabilized cells in the absence of the cellular extract or in the presence of Nap1 or Asf1, and the incorporation was analyzed by immunostaining (Figure 1-figure supplement 1C). Without the cellular extract or the histone chaperone, the exogenously added H3.1 was not detected in the permeabilized cells, indicating that no H3.1 incorporation had occurred. Nap1 promoted the promiscuous incorporation irrespective of DNA replication, and Asf1 facilitated H3.1 accumulation in the nucleoli. These data indicated that the functional deposition of the exogenously added histone complex requires the cellular extract, which may contain essential components. We conclude that the RhIP assay reproduces cellular histone deposition, and is suitable for analyzing histone incorporation *in vitro*.

### H2A.Z incorporation into chromatin differs from that of H2A and H2A.X

Among the H2A family members, canonical H2A and the H2A.X variant show even and broad genome-wide distributions, but H2A.Z specifically localizes in open chromatin, including promoters and enhancers (Buschbeck & Hake, 2017; Raisner et al., 2005). To test whether this difference reflects their deposition manners, we performed the RhIP assay (Figure 2A). The H2A-H2B and H2A.Z-H2B complexes were reconstituted *in vitro* using recombinant proteins (Figure 2B) and added to permeabilized cells, which were then immunostained (Figure 2C). The H2A and H2A.Z signals were both observed in the Cy5-negative and -positive permeabilized cells, indicating that their incorporations occur irrespective of DNA replication. We also found that H2A forms foci in the S phase nuclei. We then merged the images of the H2A and Cy5 signals. The replication foci change as cells progress through S phase (Leonhardt et al., 2000). In early S phase, the replication foci are present throughout the nucleoplasm, except for the nucleoli. The foci then accumulate at the nuclear periphery and around the nucleoli. In late S phase, the foci increase in size but decrease in number. We found that the H2A signals overlapped well with the replication foci throughout replication (Figure 2C and D). The H2A.Z signals overlapped with the early replication foci to some extent, but they were clearly eliminated from the late replication foci (Figure 2C and D). In contrast to the difference between H2A and H2A.Z, the H2A and H2A.X signals overlapped well with each other, suggesting that their incorporation mechanism is the same (Figure 2-figure supplement 1). These results indicate that H2A, H2A.X, and H2A.Z can be incorporated into chromatin in a replication-independent manner; however, during S phase, H2A and H2A.X are preferentially incorporated into the chromatin of ongoing replication sites, in contrast to H2A.Z.

**Figure 2.**
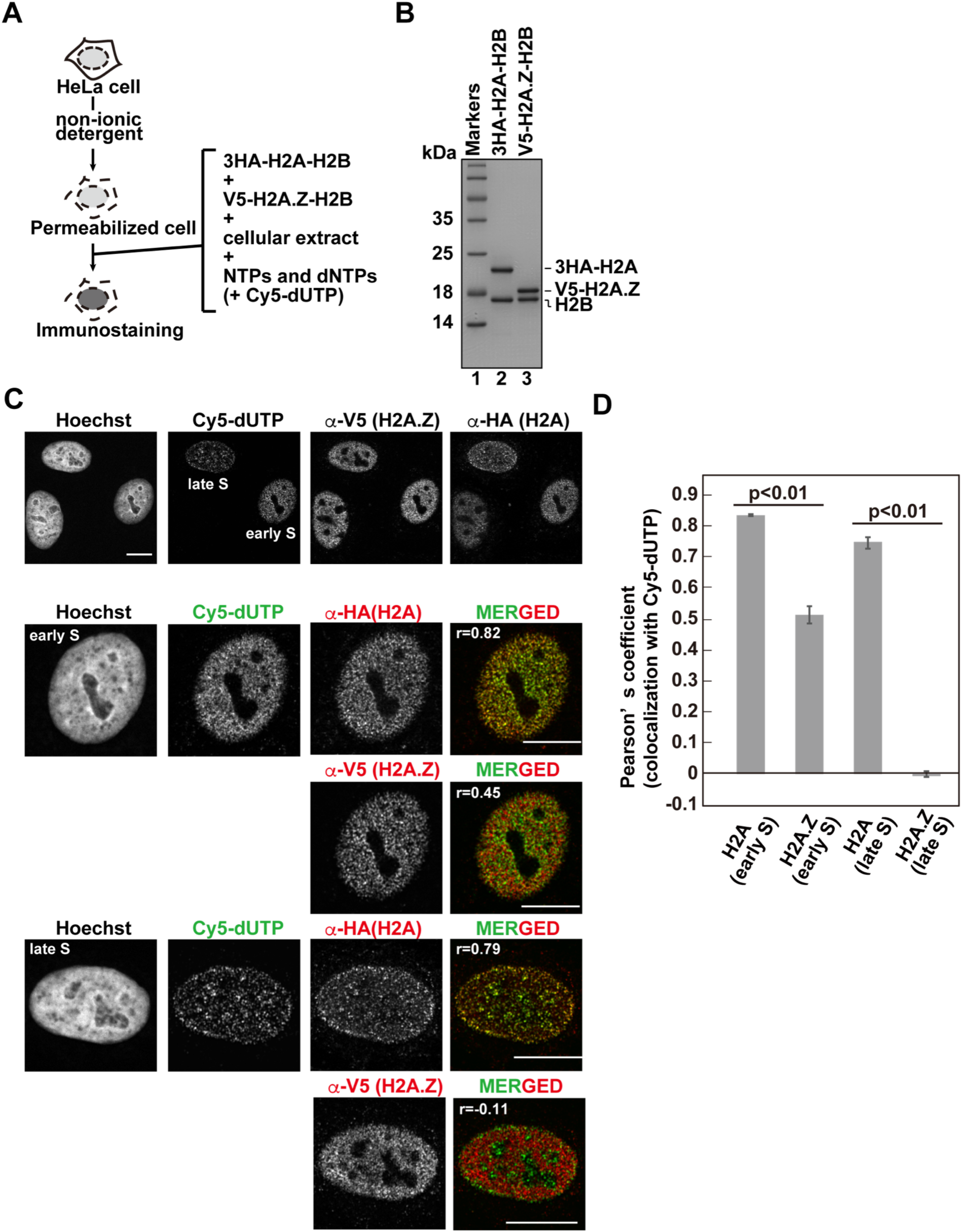
Analysis of histone H2A and H2A.Z incorporations with the RhIP assay. (A) Schematic representation of the RhIP assay, using the reconstituted H2A-H2B and H2A.Z-H2B complexes. (B) The reconstituted H2A-H2B and H2A.Z-H2B complexes were analyzed by SDS-16% PAGE with Coomassie Brilliant Blue staining. The 3HA and V5 tags were fused to the N-termini of H2A and H2A.Z, respectively. Lane 1 indicates the molecular mass markers, and lanes 2 and 3 indicate the H2A-H2B and H2A.Z-H2B complexes, respectively. (C) RhIP-immunostaining of H2A and H2A.Z. Top: Exogenously added H2A-H2B or H2A.Z-H2B complexes were stained with either an anti-HA or -V5 antibody. Cells in S phase were monitored with Cy5-dUTP. Middle and Bottom: merged images of Cy5-dUTP (green) and H2A or H2A.Z (red) in early S (Middle) and late S (Bottom) phase. Bar indicates 10 μm, and r indicates the Pearson’s correlation coefficient. (D) Colocalization analyses of Cy5-dUTP and H2A.Z or H2A (n>35 cells). Experiments were repeated three times and averaged data are shown. The two-tailed Student’s t-test was used for the statistical comparisons.

### H2A is preferentially incorporated over H2A.Z into replicating chromatin

We further analyzed the incorporation of H2A and H2A.Z into replicating chromatin by RhIP, followed by a chromatin immunoprecipitation (RhIP-ChIP) assay (Figure 3A). The reconstituted 3HA-H2A-H2B or 3HA-H2A.Z-H2B complex was added to permeabilized cells with the cellular extract and nucleotides, including Cy5-dUTP to label the nascent DNA (Figure 3A and B). After the reaction, the chromatin was partially digested by micrococcal nuclease (MNase), and the nucleosomes containing 3HA-H2A or 3HA-H2A.Z were immunoprecipitated with an antibody against the HA tag. The precipitated DNA was then extracted and analyzed by gel electrophoresis. As shown in Figure 3C, the amounts of precipitated DNA are nearly the same between the H2A and H2A.Z precipitants, as judged from the SYBR Gold staining (Figure 3C, upper); however, the amount of nascent DNA labeled with Cy5 is much greater in the H2A precipitant than in the H2A.Z sample (Figure 3C, lower). This result indicates that H2A is incorporated into replicating chromatin more efficiently than H2A.Z.

**Figure 3.**
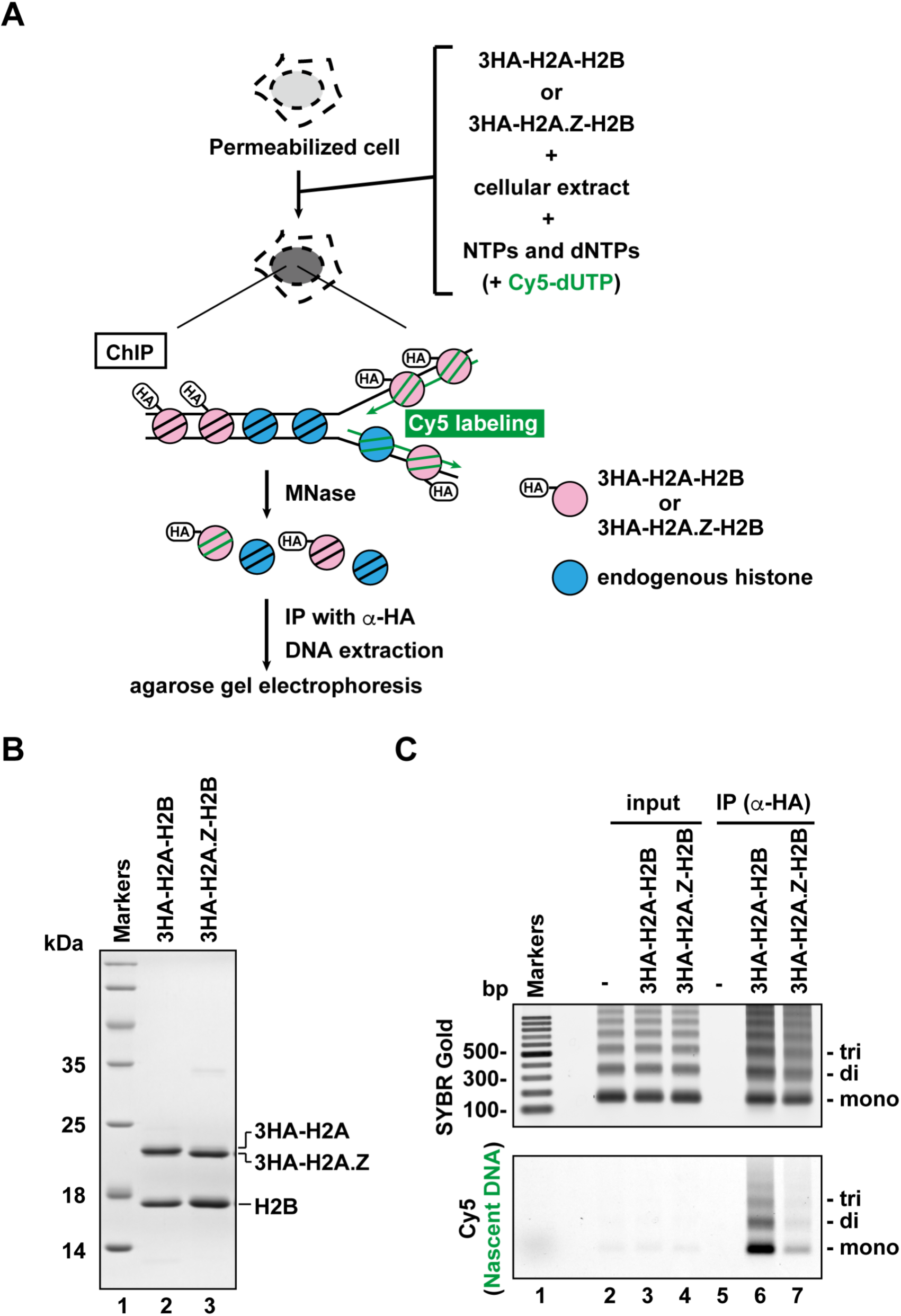
Analysis of histone H2A and H2A.Z incorporations with the RhIP-ChIP assay. (A) Schematic representation of the RhIP-ChIP assay, using the reconstituted H2A-H2B and H2A.Z-H2B complexes. The reconstituted H2A-H2B or H2A.Z-H2B complex was added to permeabilized cells with the cellular extract and nucleotides. Cy5-dUTP was added to label the nascent DNA. The chromatin was partially digested with micrococcal nuclease (MNase). Chromatin immunoprecipitation was performed with anti-HA magnetic beads. The precipitated DNA was extracted and analyzed by agarose gel electrophoresis. (B) Reconstituted H2A-H2B and H2A.Z-H2B complexes were analyzed by SDS-16% PAGE with Coomassie Brilliant Blue staining. A 3HA tag was fused to the N-termini of H2A and HA.Z. Lane 1 indicates the molecular mass markers, and lanes 2 and 3 indicate the H2A-H2B and H2A.Z-H2B complexes, respectively. (C) The immunoprecipitated DNA was analyzed by 2% agarose electrophoresis. Upper and lower images were obtained from the same gel. The DNA was visualized with SYBR Gold (upper), and the nascent DNA was visualized by detecting the Cy5 signals (lower). Lane 1 indicates the 100 bp DNA ladder. Lanes 2-4 and 5-7 indicate input samples and immunoprecipitated samples, respectively. Each lane indicates a negative control without the reconstituted histone complex (lanes 2 and 5), the samples with 3HA-H2A-H2B (lanes 3 and 6), and the samples with 3HA-H2A.Z-H2B (lanes 4 and 7).

The replication timing in S phase strongly correlates with the chromatin configurations (Rivera-Mulia & Gilbert, 2016). In general, early and late replicating chromatin regions correspond to open and closed chromatin, respectively. We investigated whether the efficiencies of H2A and H2A.Z incorporation into replicating chromatin change, according to the replication timing (Figure 3-figure supplement 1). For this analysis, the cells were synchronized in early S phase by a double thymidine block, and then early and late S phase cells were collected at 0 and 5 hours post thymidine-release, respectively. The synchronized cells showed the typical early and late replication foci representing nascent DNA labeled with Cy5-dUTP (Figure 3-figure supplement 1A). Using these cells, we performed RhIP-ChIP assays of H2A and H2A.Z. The results revealed that the incorporation efficiencies of both H2A and H2A.Z into replicating chromatin did not change, irrespective of the replication timing in S phase. This implies that the efficiencies of replication-coupled histone deposition are not different between open (early S-replicating) and closed (late S-replicating) chromatin (Figure 3-figure supplement 1B).

### Open and closed chromatin structures regulate histone deposition

The RhIP-ChIP assay showed that the replication-coupled deposition of H2A.Z was constant in the early and late S phases (Figure 3C and Figure 3-figure supplement 1). In contrast, the RhIP-immunostaining revealed that more signals of H2A.Z incorporation were overlapped at the early replicating foci than the late replicating foci (Figure 2C and D). This discrepancy may arise from the lower resolution of the immunostaining imaging. Some of the overlapping signals of H2A.Z and Cy5 in early S phase might represent the replication-independent H2A.Z deposition that occurred close to, but not exactly at, the replication sites in open chromatin. To determine whether the efficiency of the histone deposition depends on the open/closed chromatin configuration, we performed RhIP-ChIP-seq and analyzed the H2A and H2A.Z incorporations in each type of chromatin (Figure 4). First, we investigated the replication-independent histone deposition using asynchronous permeabilized cells, in which the majority of the cells are out of S phase. We found that the RhIP-ChIP-seq profiles of H2A and H2A.Z showed specific peaks at megabase resolution, which were not observed in the input samples (Figure 4A upper). We noticed that these patterns are similar to the DNaseI-seq results, which mapped open chromatin regions (Tsompana & Buck, 2014), suggesting that H2A and H2A.Z are predominantly incorporated into open chromatin regions. Furthermore, by comparing each profile at the kilobase resolution, we found that H2A.Z predominantly accumulates at the transcription start sites (TSS) of open chromatin regions (Figure 4A lower and 4B). We then analyzed the efficiency of histone incorporations into open and closed chromatin using the chromHMM data (Core 15-state model), which classified the open and closed chromatin regions (Ernst & Kellis, 2012; Roadmap Epigenomics Consortium et al., 2015) (Figure 4C and Figure 4-figure supplement 1A). The ChIP/input ratios of each chromatin region revealed that H2A and H2A.Z were efficiently incorporated into transcriptionally active open chromatin, while their incorporations into closed chromatin were inefficient. These results indicate that histone incorporation occurs mainly in the open chromatin regions in a replication-independent manner, and closed chromatin suppresses histone incorporation.

**Figure 4.**
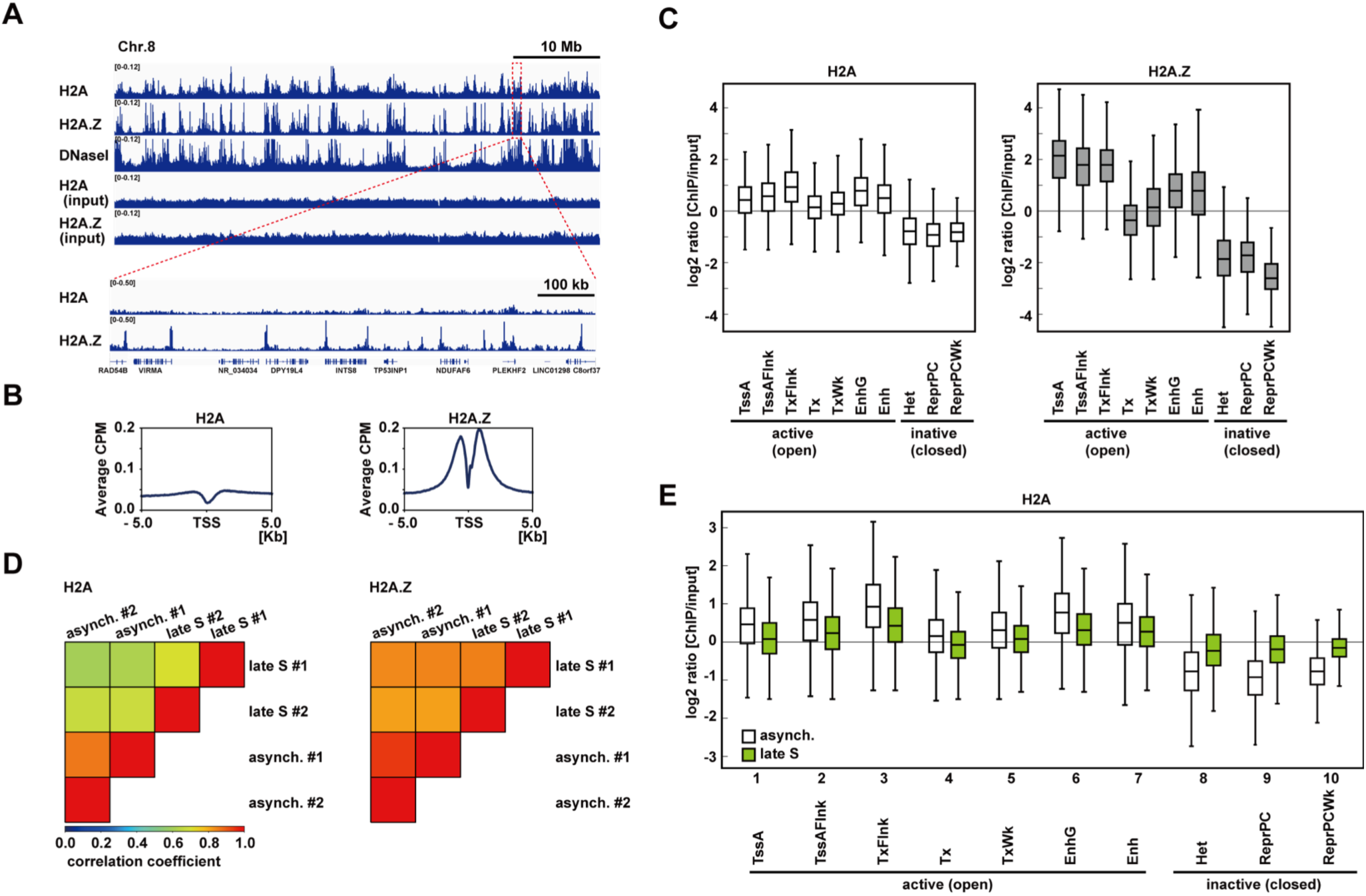
RhIP-ChIP-seq analysis of H2A and H2A.Z. (A) RhIP-ChIP-seq and DNaseI-seq profiles using asynchronous cells were visualized with the Integrative Genomics Viewer. From top to bottom, profiles of H2A, H2A.Z, DNaseI-seq (GEO:GSM816643), input (H2A), and input (H2A.Z) are indicated. (B) Aggregation plots of the H2A (left), and H2A.Z (right) localizations around all TSSs. (C) Enrichment of incorporated H2A (left) or H2A.Z (right) in asynchronous cells. Each chromatin region was previously annotated by the chromHMM, as follows. TssA: active TSS, TssAFlnk: flanking active TSS, TxFlnk: transcribed state at the 5’ and 3’ ends of genes showing both the promoter and enhancer signatures, Tx: strong transcription, TxWk: weak transcription, EnhG: genic enhancers, Enh: enhancers, Het: heterochromatin, ReprPC: repressed PolyComb, and ReprPCWk: weak repressed PolyComb (Ernst & Kellis, 2012; Roadmap Epigenomics Consortium et al., 2015). (D) The Pearson’s correlation coefficients between asynchronous and late S cells (two biological replicates each) were calculated from the RhIP-ChIP-seq data at 800 bp intervals. Left and right panels indicate the correlation coefficients of the H2A and H2A.Z data, respectively. (E) Enrichment of incorporated H2A in asynchronous (white boxes) and late S (green boxes) phase cells. Data for asynchronous cells are those shown in Figure 4C.

We then examined whether the efficiency of the histone incorporation into closed chromatin changes during S phase (Figure 4D and E). We performed RhIP-ChIP-seq with cells synchronized at late S phase. We found that the RhIP-ChIP-seq profiles between asynchronous cells and late S phase cells changed for H2A, but not for H2A.Z, indicating that H2A incorporation into chromatin is affected by replication, in contrast to H2A.Z incorporation (Figure 4D). We then investigated how the incorporation efficiency of H2A into open and closed chromatin regions changes between asynchronous and late S phase cells. The results revealed that the efficiency of H2A incorporation into open chromatin decreased in late S phase (Figure 4E, lanes 1-7), while in contrast, the incorporation into closed chromatin increased (Figure 4E, lanes 8-10 and Figure 4-figure supplement 1B). Considering the fact that only closed chromatin is replicated in late S phase, the changes in the incorporation efficiency may be due to the changes in replicating chromatin. Note that replication does not progress completely in the RhIP assay, and only small fraction of closed chromatin is replicated under the conditions shown in Figure 4E, which is the reason why the ChIP/input ratio does not exceed one. We concluded that replication is required for the incorporation of H2A into closed chromatin. This indicates that, in principle, H2A.Z-H2B incorporation into chromatin is not coupled with replication, which leads to H2A.Z elimination from closed chromatin.

### The αC helix of H2A is important for its replication-coupled deposition

As the RhIP assay reproduces the replication-coupled and replication-independent incorporations of H2A and H2A.Z, we then tried to identify the residues responsible for the replication-coupled H2A and replication-independent H2A.Z depositions by a mutant analysis. A previous study showed that swapping the M6 region of H2A.Z with the corresponding H2A residues could not rescue the embryonic lethality of the H2A.Z null mutation in *Drosophila melanogaster* (Clarkson et al., 1999), suggesting that the region specifies the H2A.Z identity. The M6 region of H2A.Z and the corresponding region of H2A are exposed on the surface of the H2A.Z-H2B or H2A-H2B dimer (Horikoshi et al., 2013; Luger et al., 1997; Suto, Clarkson, Tremethick, & Luger, 2000; Tachiwana et al., 2010) (Figure 5A, cyan or green). This indicates that another protein can recognize the regions, which may be important for their depositions. To test this idea, we constructed the swapped mutant (H2A.Z_M6) and performed the RhIP assay (Figure 5B-F). Surprisingly, the H2A.Z_M6-H2B signals were observed in late replicating chromatin (Figure 5D and E), and its incorporation pattern in late S phase was more similar to that of H2A-H2B, rather than H2A.Z-H2B (Figure 5F). This indicated that the mutant is no longer H2A.Z, in terms of deposition. Thus, the M6 region of H2A.Z is responsible for the H2A.Z-specific deposition, and the corresponding region (amino acids 89-100) of H2A is responsible for the replication-coupled H2A deposition.

**Figure 5.**
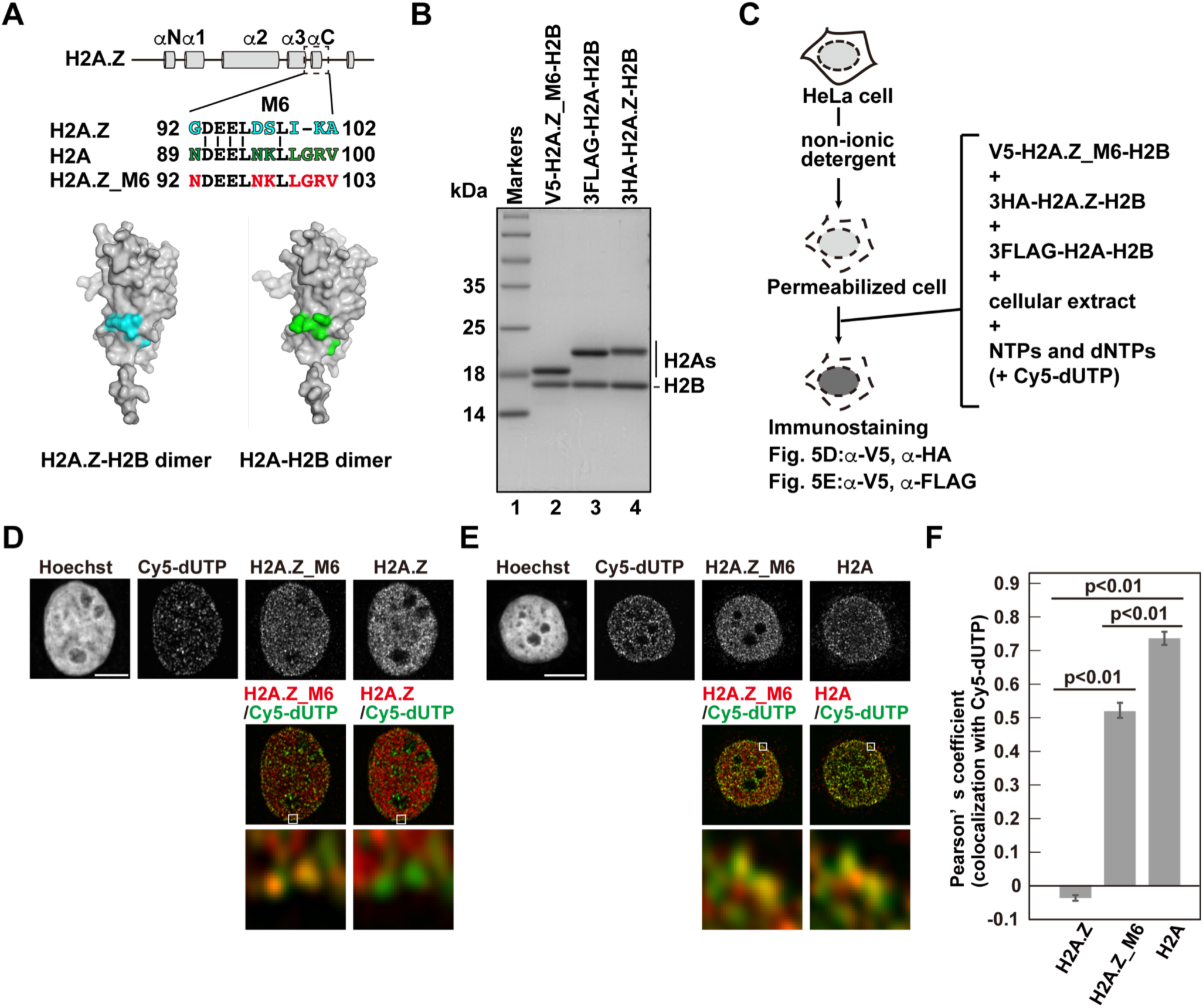
Identification of responsible residues for H2A- and H2A.Z-specific incorporations. (A) Amino acid alignments of the H2A.Z M6 region and its counterpart in H2A (upper). The structural models of the H2A.Z-H2B and H2A-H2B dimers (PDB IDs: 3WA9 and 3AFA, respectively). The specific residues are highlighted in cyan or green, respectively. All residues are located on the surface of the dimers. (B) Reconstituted H2AZ_M6-H2B, H2A-H2B, and H2A.Z-H2B complexes were analyzed by SDS-16% PAGE with Coomassie Brilliant Blue staining. H2AZ_M6, H2A, and H2A.Z were expressed as N-terminally V5, 3FLAG, and 3HA fused proteins, respectively. Lane 1 indicates the molecular mass markers, and lanes 2-4 indicate the H2A.Z_M6-H2B, H2A-H2B, and H2A.Z-H2B complexes, respectively. (C) Schematic representation of the RhIP assay, using the reconstituted H2A.Z_M6-H2B, H2A-H2B, and H2A.Z-H2B complexes. (D) RhIP-immunostaining images of H2A.Z and H2A.Z_M6. Exogenously added H2A.Z-H2B and H2A.Z_M6-H2B complexes were stained with the anti-V5 or -HA antibody. Cells in S phase were monitored with Cy5-dUTP. Middle: merged images of Cy5-dUTP (green) and H2A.Z_M6 (red) or H2A.Z (red). Bottom: magnified images of boxed areas are shown. Bar indicates 5 μm. (E) RhIP-immunostaining images of H2A and H2A.Z_M6. Exogenously added H2A-H2B and H2A.Z_M6-H2B complexes were stained with the anti-V5 or -FLAG antibody. Cells in S phase were monitored with Cy5-dUTP. Middle: merged images of Cy5-dUTP (green) and H2A.Z_M6 (red) or H2A (red). Bottom: magnified images of boxed areas are shown. Bar indicates 5 μm. (F) Colocalization analyses of Cy5-dUTP and H2A.Z, H2A.Z_M6 or H2A in late S phase (n>35 cells). Experiments were repeated three times and averaged data are shown. The two-tailed Student’s t-test was used for the statistical comparisons.

## Discussion

We established the novel RhIP assay, combining permeabilized cells and reconstituted histone complexes, to analyze histone incorporation. Previous approaches using genetically encoded histone genes have revealed chromatin dynamics, including nucleosome turnover kinetics, but have limitations on the time resolution, as they require time to synthesize and/or label histones (Deal & Henikoff, 2010). In contrast, the RhIP assay can detect histone incorporation with better time resolution, as it does not require histone synthesis or labeling. In fact, we could analyze histone incorporations in the early and late S phases separately, using synchronized cells (Figure 3-figure supplement 1 and Figure 4E). By combining ChIP-seq, RhIP assay enables to analyze histone incorporation at the DNA sequence level, while ChIP-seq reveals their static presence. In addition, the RhIP-ChIP-seq analysis of H2A.Z shows better enrichment (S/N ratio) as compared with ChIP-seq (Figure 4-figure supplement 2), probably due to the high affinity antibody against the epitope-tag. Together with the fact that the RhIP-ChIP assay requires a small number of cells (approximately 8*×*10^6^ cells), the RhIP assay is suitable for the biochemical analysis of ChIP samples. The effects of post-translational modifications (PTMs) of histones on their incorporation have remained elusive. Methods to produce histones with PTMs *in vitro* have been developed (Nadal, Raj, Mohammed, & Davis, 2018), and thus the analysis of the effects of PTMs on histone incorporation will be the next target for the RhIP assay.

The present study has revealed the correlation between histone incorporation and chromatin structure, using the RhIP assay. We analyzed the incorporation of the H2A family members, H2A, H2A.X, and H2A.Z. We found that the incorporations of H2A-H2B and H2A.X-H2B into transcriptionally active open chromatin can occur in both replication-coupled and -independent manners, whereas their incorporation into transcriptionally inactive closed chromatin occurs only in a replication-coupled manner in late S phase (Figures 4 and 6). This indicated that histone exchange would rarely occur in closed chromatin. Thus, the genome-wide localization of H2A and H2A.X is achieved by a combination of deposition into open chromatin and replication-coupled deposition into closed chromatin (Figure 6). In contrast, H2A.Z exhibited a much lower frequency of replication-coupled deposition, as compared to H2A (Figure 3C and Figure 3-figure supplement 1). Together with the fact that the amount of H2A.Z is much lower than that of H2A in S phase cells (Wu et al., 1982), we concluded that little to no H2A.Z is incorporated into closed chromatin (Figure 6). This is consistent with previous observations that the H2A.Z and DNA methylation localizations are mutually exclusive (Nothjunge et al., 2017; Zilberman, Coleman-Derr, Ballinger, & Henikoff, 2008), and that H2A.Z predominantly exists at promoters and enhancers (Buschbeck & Hake, 2017; Raisner et al., 2005). The means by which H2A.Z becomes enriched at specific regions of open chromatin have remained enigmatic. Our study suggested that the H2A.Z elimination from transcriptionally inactive chromatin is due to the low frequencies of histone exchange and replication-coupled H2A.Z incorporation, which may partially explain the H2A.Z distribution pattern (Figure 6). Moreover, H2A.Z was specifically incorporated into chromatin around the TSS in the RhIP assay, in excellent agreement with the steady state localization of H2A.Z revealed by ChIP-seq (ENCODE Project Consortium, 2012). There may be a specific mechanism for H2A.Z targeting to the TSS. H2A.Z is also found at pericentric heterochromatin (Boyarchuk et al., 2014; Greaves et al., 2007; Rangasamy et al., 2003). As human pericentric heterochromatin is composed of repetitive DNA sequences (satellite repeats), we could not analyze H2A.Z incorporation into pericentric heterochromatin by the RhIP-ChIP-seq assay (Figure 4). Previous studies showed that the transcriptional activation of pericentric satellites accompanies the structural changes from heterochromatin to euchromatin (Jolly et al., 2004; Rizzi et al., 2004; Saksouk, Simboeck, & Déjardin, 2015; Valgardsdottir et al., 2005). In addition, the pericentric region of human chromosome 9 is heterogeneous, with both heterochromatic and euchromatic regions (Gilbert et al., 2004). Therefore, H2A.Z incorporation into pericentric heterochromatin may occur, as in the open chromatin of the region.

**Figure 6.**
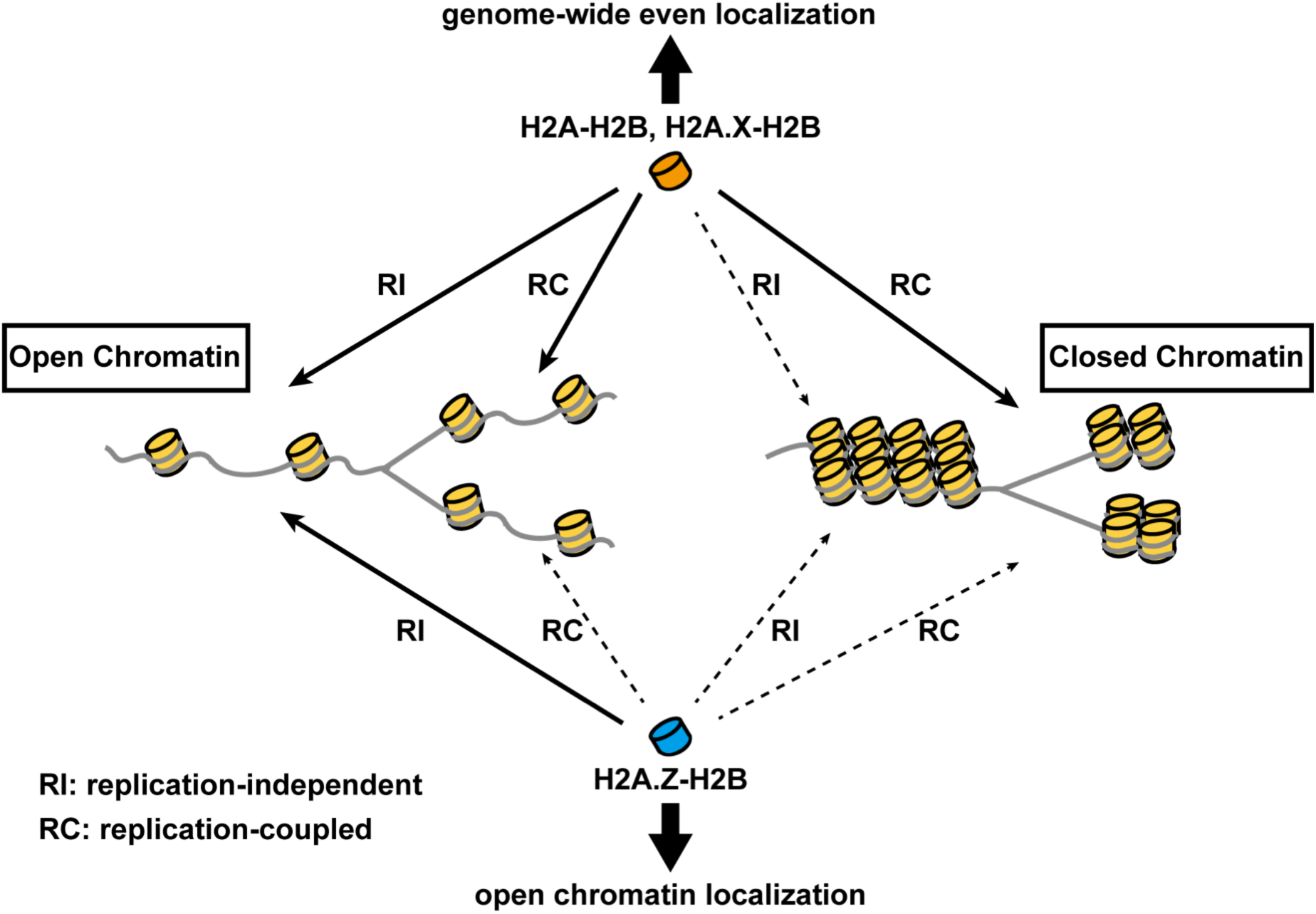
Model of differential histone incorporations into open and closed chromatin. In open chromatin, the H2A-H2B and H2A.X-H2B complexes are incorporated in replication-independent (RI) and replication-coupled (RC) manners, while H2A.Z-H2B is incorporated only in a replication-independent manner. In closed chromatin, new histone depositions of H2A and H2A.X, but not H2A.Z, occur only in a replication-coupled manner. This leads to the global localizations of H2A-H2B and H2A.X-H2B. It also leads to the specific localization of H2A.Z-H2B, including its elimination from closed chromatin.

Another H2A variant, MacroH2A, localizes in closed chromatin. A recent study showed that the *de novo* incorporation of MacroH2A is genome-wide. It is then removed from transcribed chromatin in a transcription-associated mechanism, leading to its specific localization at repressed chromatin (Sun et al., 2018). Our data may be able to explain why this two-step mechanism is needed for MacroH2A localization in closed chromatin. We showed that H2A is incorporated into closed chromatin in a replication-coupled manner. If MacroH2A also uses the replication-coupled deposition mechanism, then it is inevitably incorporated into closed and open chromatin, first resulting in the genome-wide distribution. MacroH2A must then be removed from the active chromatin for its specific closed chromatin localization.

We found that histone incorporations are regulated by the chromatin structure, which may be important for maintaining the closed chromatin configuration. Histones reportedly form a pre-deposition complex, which includes many transcription-related factors, before their incorporation into chromatin *in vivo* (Dunleavy et al., 2009; Mao et al., 2014; Obri et al., 2014; Tagami et al., 2004). For instance, the H2A-H2B pre-deposition complex contains Spt16 and SSRP1, which form the heterodimer complex FACT that functions in transcription facilitation, and the pre-deposition H2A.Z-H2B complex also includes a chromatin remodeling factor (SRCAP), a histone acetyltransferase (Tip60), and acetyl-lysine binding proteins (GAS41, Brd8). If histone exchange usually occurs in closed chromatin, then these transcription-related factors might accumulate in the closed chromatin and alter the epigenetic chromatin states. Therefore, the deficiency of replication-independent histone exchange in closed chromatin may be important for maintaining a transcriptionally inactive state.

In spite of the high sequence homology between H2A and H2A.Z, their localizations are different. By using the swapping mutant, we analyzed the residues responsible for the specific depositions of H2A and H2A.Z (Figure 5). We identified residues 88-100 of H2A as being responsible for its replication-coupled deposition. Although the means by which this region contributes to the incorporation into replicating chromatin remain unknown, a factor that binds this region and allows H2A to assemble at a replicating site may exist. The major H2A-specific chaperones, Spt16 and Nap1, which are components of the H2A pre-deposition complex, do not bind to this region (Aguilar-Gurrieri et al., 2016; Kemble, McCullough, Whitby, Formosa, & Hill, 2015). Thus, the protein that recognizes these residues is likely to be a chromatin protein involved in replication, rather than a component of the H2A pre-deposition complex. This region is a counterpart of the H2A.Z M6 region, and the H2A.Z_M6 swapping mutant did not compensate for the embryonic lethality of the H2A.Z knockout in *Drosophila melanogaster* (Clarkson et al., 1999). As this region is essential for the H2A.Z eviction by ANP32E (Gursoy-Yuzugullu, Ayrapetov, & Price, 2015; Obri et al., 2014), the incorporated H2A.Z_M6 mutant may not be removed from closed chromatin, thus resulting in aberrant gene expression and impaired embryonic development.

In conclusion, our novel method elucidated the mechanism of histone incorporation at the DNA sequence level, and revealed that the chromatin structure is the first determinant of histone localization.

## Materials and Methods

### Cell culture and thymidine block

HeLa cells were cultured in DMEM medium supplemented with 10% fetal bovine serum, at 37°C in a 5% CO_2_ atmosphere. For cell synchronization, HeLa cells were cultured with 2 mM thymidine for 19 hours. The medium was then changed to remove the thymidine. After 9 hours of culture without thymidine, the HeLa cells were cultured with 2 mM thymidine again for 15 hours, for synchronization in S phase.

### Reconstitution of histone complex

The human H3.1 and H3.3 genes were inserted in the pET21a vector (Novagen), as C-terminal epitope-tag-His_6_ fused genes. The human H2A, H2A.X, H2A.Z, and H2A.Z_M6 genes were inserted in the pET15b vector (Novagen) as N-terminal His6-epitope-tag fused genes. All of the genes were overexpressed in the BL21(DE3) *E. coli* strain, by adding 0.5 mM isopropyl-β-D-thiogalactopyranoside. Each histone was then purified as described previously, using Ni-NTA affinity chromatography (Tachiwana et al., 2010). The epitope tag-fused histones were freeze-dried without removing the His_6_ tags. Human H2B and H4 were overexpressed and purified after removing the His_6_ tags, as described previously (Tachiwana et al., 2010). Freeze-dried H3.1 or H3.3 was mixed with H4 and H2A, H2A.X, H2A.Z, and H2A.Z_M6, along with H2B, in 20 mM Tris-HCl buffer (pH 7.5), containing 7 M guanidine hydrochloride and 20 mM 2-mercaptoethanol, and incubated on ice. After 1 hour, the samples were dialyzed against reconstitution buffer (10 mM Tris-HCl, pH 7.5, and 2 mM 2-mercaptoethanol) containing 2 M NaCl, overnight at 4°C. The NaCl concentration was then decreased by three steps of dialysis against reconstitution buffer containing 1 M NaCl for 4 hours, 0.5 M NaCl for 4 hours, and 0.1 M NaCl overnight. After the dialyses, the precipitants were removed by centrifugation and the supernatants were analyzed by Superdex 200 gel filtration chromatography (GE Healthcare).

### RhIP assay

Cell permeabilization was performed as described previously, with minor modifications (Kimura et al., 2006). HeLa cells were chilled on ice and rinsed twice with ice-cold PBF (100 mM CH_3_COOK, 10 mM Na_2_HPO_4_, 30 mM KCl, 1 mM dithiothreitol, 1 mM MgCl_2_, 1 mM ATP, and 5% Ficoll). Afterwards, PBF containing 0.2% Triton X-100 was added to the cells. After a 5 min incubation on ice, the permeabilized cells were rinsed twice with ice-cold PBF. The preparation of the cellular extract was described previously, as a cytosolic extract (Martini, Roche, Marheineke, Verreault, & Almouzni, 1998). The histone incorporation assay was performed as described previously, with modifications (Kimura et al., 2006). A reaction mixture, containing 100 nM histone complex, 60% cellular extract, 2.5% Ficoll, 100 mM CH_3_COOK, 10 mM Na_2_HPO_4_, 30 mM KCl, 1 mM dithiothreitol, 1 mM MgCl_2_, 100 μM each of dNTPs, and NTPs (Roche) with or without 250 nM Cy5-dUTP (Enzo Life Sciences), was added to the permeabilized cells. After 60 min at 30°C, the cells were washed twice with PBS containing 0.05% Tween 20 (PBST), for 5 min at room temperature.

### RhIP-immunostaining

After the RhIP assay, the cells were fixed with 4% PFA (Electron Microscopy Sciences) in PBS for 20 min at room temperature, and rinsed three times with PBS. After fixation, the cells were treated with 1% BSA in PBST for 1 hour at room temperature, and then with the primary antibodies in the same buffer. After 2 hours at room temperature, the cells were washed with PBST three times for 10 min, treated with the secondary antibodies for 1 hour at room temperature, and washed with PBST three times for 10 min. DNA was stained with Hoechst 33342. The samples were mounted with ProLong Gold (Life Technologies). The images in Figure 1, Figure 1-figure supplement 1, Figure 2-figure supplement 1, and Figure 3-figure supplement 1 were acquired by using the Deltavision set-up (GE Healthcare) with an inverted Olympus IX71 microscope, equipped with a CoolSNAP ES2 CCD camera (Photometrics) and a 60×, 1.42 Plan Apo N Olympus oil-immersion objective. Other images were acquired with an LSM 880 inverted confocal microscope (Zeiss), equipped with an AiryScan module and a 63×, 1.40 Plan-Apochromat Zeiss oil objective. All image files were converted to the TIFF format using the ImageJ software (Schneider, Rasband, & Eliceiri, 2012) and imported into Illustrator (Adobe) for assembly. The co-localization analysis was performed using the ImageJ Colocalization_Finder plugin.

### RhIP-ChIP

After the RhIP assay, the chromatin was digested with micrococcal nuclease (MNase) as described previously (Tachiwana et al., 2015). The reaction was terminated by adding 10 mM EDTA, and the solubilized chromatin fragments were separated from the pellets by centrifugation at 16,000g for 5 min at 4°C. The samples were then mixed with 50 μl of anti-HA-tag mAb-Magnetic Beads (MBL International) in 15 mM Tris-HCl, pH 7.5, 300 mM NaCl, and 0.1% NP-40. After an overnight incubation at 4°C with gentle mixing on a wheel, the beads were washed twice with 1 ml of wash buffer (10 mM Tris-HCl, pH 7.5, 2 mM EDTA, 500 mM NaCl, and 0.1% Triton X-100). The DNA was then eluted with a Proteinase K solution (20 mM Tris-HCl, pH 8.0, 20 mM EDTA, 0.5% sodium dodecyl sulfate (SDS), and 0.5 mg/ml Proteinase K) and extracted with phenol-chloroform. The DNA was precipitated with ethanol and resuspended in 10 mM Tris-HCl, pH 8.0, and 1 mM EDTA. The resulting DNA samples were separated by electrophoresis on a 2% agarose gel (7.4 V/cm, 35 min) or a 1.5 % agarose gel (7.1 V/cm, 60 min) in 1×TAE, and stained with SYBR Gold (Thermo Fisher). Images of the DNA stained with SYBR Gold and the nascent DNA labeled with Cy5 were obtained with an Amersham Typhoon scanner (GE Healthcare).

### RhIP-ChIP-seq

After the RhIP assay, the permeabilized cells were treated with 3% formaldehyde in PBS for 5 min at room temperature. The fixed chromatin of the permeabilized cells was digested with MNase in ChIP buffer (10 mM Tris-HCl, pH 8.0, 200 mM KCl, 1 mM CaCl_2_, 0.5% NP-40). The supernatant was incubated with 50 μl of anti-HA-tag mAb-Magnetic Beads, and gently mixed by rotation at 4°C overnight. The beads were then washed three times with ChIP buffer containing 500 mM KCl and TE. After the beads were washed, they were resuspended in 100 μl of ChIP elution buffer (50 mM Tris-HCl, pH 8.0, 10 mM EDTA, and 1% SDS) containing 25 mM NaCl, and incubated overnight at 65°C to reverse the cross-links. The DNA was then eluted with 0.4 mg/ml Proteinase K. The eluted DNA samples were further purified with a NucleoSpin Gel and PCR clean-up kit (MACHEREY-NAGEL). The DNA libraries were prepared using a SMARTer ThruPLEX Tag-seq Kit (Takara Bio), and the samples were sequenced on an Illumina HiSeq1500 system.

### ChIP-Seq data analysis

The in-read unique-molecular-identifiers (tags) were extracted using UMI-tools (Smith, Heger, & Sudbery, 2017) with the command: umi_tools extract --extract-method=regex --bc-pattern=’(?P<UMI_1>.{6})(?P<DISCARD_1>.{0,3}GTAGCTCA){s<=2}’. The extracted reads were mapped to the human genome (GRCh38) using hisat2 (version 2.1.0) (Kim, Langmead, & Salzberg, 2015). The PCR-duplicates were removed using umi_tools dedup. The read counts on 15 chromatin states were calculated using BED-Tools (Quinlan & Hall, 2010). The definition of the ChromHMM track was obtained from the consolidated data set of the Roadmap Epigenomics project (E117_15_coreMarks_hg38lift_mnemonics.bed) (Roadmap Epigenomics Consortium et al., 2015). The overall concentrations of the ChIP signals (log2 ratio) were calculated as the ratio of the proportion of reads in each chromatin state between the ChIP and input DNA data; *i.e*., log_2_(ChIP/Input) after normalization of the total reads. The signal tracks (bigwig files) were created by 1 bp intervals on the genome, and then the counts were normalized as CPM (Reads Per Million reads), using deepTools (Ramírez, Dündar, Diehl, Grüning, & Manke, 2014). ChIP-seq signals were visualized with the Integrative Genomics Viewer, IGV (Robinson et al., 2011). The Pearson correlation coefficients were calculated throughout 800 bp intervals with the multiBigwigSummary program of deepTools, and plotted with the plotCorrelation program (Ramírez et al., 2014). The computeMatrix and plotFingerprint programs of deepTools were utilized to analyze the percentages of peak localizations around the TSS and the enrichment of ChIP signals, respectively.

### Purification of Nap1 and Asf1

Human Nap1 was purified as described previously (Tachiwana et al., 2008). The human Asf1 gene was inserted in the pET15b vector (Novagen), in which the thrombin proteinase recognition sequence was replaced by the PreScission protease recognition sequence. *Escherichia coli* strain BL21-CodonPlus (DE3)-RIL cells (Agilent Technologies) were freshly transformed with the vector, and cultured at 30°C. After the cell density reached an *A*_600_ = 0.8, 1 mM isopropyl-β-D-thiogalactopyranoside was added, and the culture was continued at 18°C for 12 h to induce His_6_-tagged Asf1 expression. The cells were collected and resuspended in 20 mM Tris-HCl, pH 7.5, containing 500 mM KCl, 10% glycerol, 0.1% NP-40, 1 mM phenylmethylsulfonyl fluoride (PMSF), and 2 mM 2-mercaptoethanol. After cell disruption by sonication, the debris was removed by centrifugation (27,216 *×*g; 20 min), and the clarified lysate was mixed gently with 4 ml (50% slurry) of nickel-nitrilotriacetic acid (Ni-NTA)-agarose resin (Qiagen) at 4°C for 1 h. The Ni-NTA beads were washed with 200 ml of 20 mM Tris-HCl, pH 7.5, containing 500 mM NaCl, 10% glycerol, 10 mM imidazole, and 2 mM 2-mercaptoethanol. The His_6_-tagged Asf1 was eluted by a 100-ml linear gradient of 10 to 500 mM imidazole in 50 mM Tris-HCl buffer (pH 7.5), containing 500 mM NaCl, 10% glycerol, and 2 mM 2-mercaptoethanol. PreScission protease (8 units/mg protein, GE Healthcare) was added to remove the His_6_ tag from the Asf1. The sample was dialyzed against 20 mM Tris-HCl, pH 7.5, containing 100 mM NaCl, 1 mM EDTA, 10% glycerol, and 2 mM 2-mercaptoethanol. The Asf1 was further purified by chromatography on a Mono Q (GE Healthcare) column, eluted with a 25-ml linear gradient of 100–600 mM NaCl in 20 mM Tris-HCl, pH 7.5, containing 1 mM EDTA, 10% glycerol, and 2 mM 2-mercaptoethanol. The eluted Asf1 was further purified by chromatography on a Superdex 75 (GE Healthcare) column, eluted with 1.2 column volumes of the same buffer containing 100 mM NaCl. The Asf1 was repurified by Mono Q chromatography, concentrated, and dialyzed against 20 mM Tris-HCl (pH 7.5), containing 150 mM NaCl, 1 mM dithiothreitol, 0.5 mM EDTA, 0.1 mM PMSF, and 10% glycerol.

### Antibodies

For immunostaining, anti-HA (mouse, 1/1,000, Santa Cruz sc-7392), anti-DDDDK (FLAG) (rabbit, 1/500, MBL, PM020), or anti-V5 (chicken, 1/1,000, Abcam, ab9113) was used as the primary antibody. Goat Alexa Fluor 488 or 546-conjugated anti-mouse IgG (Life Technologies), or goat DyLight 488 or 550-conjugated anti-rabbit IgG or chicken IgY (Thermo Fisher) was used as the secondary antibody.

### Data Availability

The deep sequencing data in this study are publicly available at the accession number GEO: GSE130947.

## Acknowledgments

This work was supported by JSPS KAKENHI Grant Numbers JP17H05013, JP16K14785 (to H.T.), JP18H05531, JP18K19310 (to N.S.), JP18H04904, JP19H04970, JP19H03158 (to K.M.), JP18K19432, JP19H05425, JP19H03211 (to A.H.), and JP17K19356, JP17H03608, JP18H05527, JP18H04802, JP19H05244 (to Y.O.), and JST CREST grants (JPMJCR16G1 to Y.O., H.Kurumizaka, and H.Kimura), JP18H05534 and JP17H01408 (to H.Kurumizaka), and JP17H01417 and JP18H05527 (to H.Kimura). We thank Dr. Yuma Ito at Tokyo Institute of Technology for technical advice. We also thank Dr. Crawford at Duke University for the DNaseI-seq data (GEO:GSM816643) and the ENCODE Consortium. H.T. is supported by The Nakajima Foundation. N.S. is supported by the Naito Foundation, the Takeda Science Foundation, and the Sharyo Foundation.

## Competing interests

All authors have and declare that: (i) no support, financial or otherwise, has been received from any organization that may have an interest in the submitted work; and (ii) there are no other relationships or activities that could appear to have influenced the submitted work.

---

**Figure 1- figure supplement 1.**
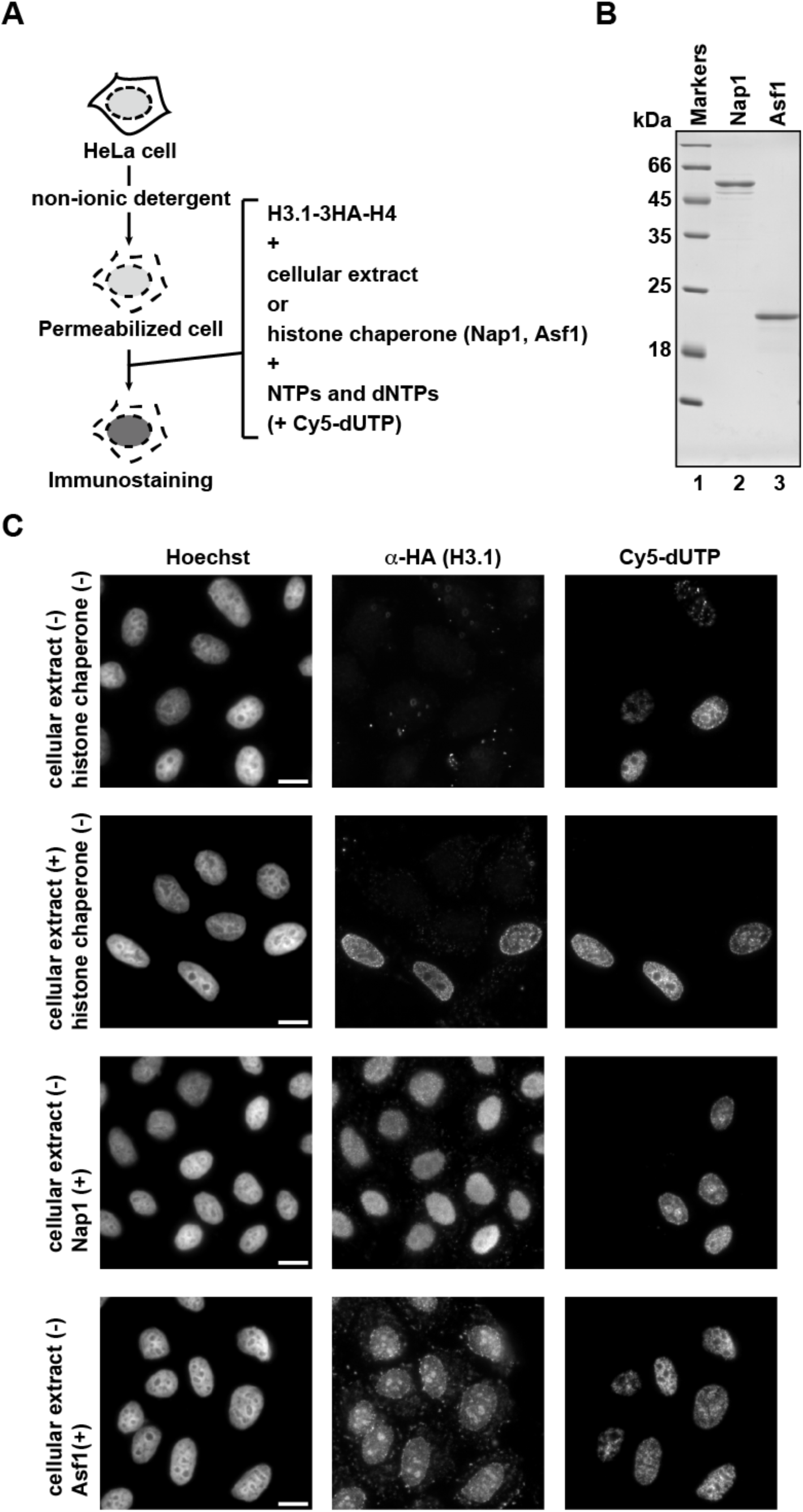
RhIP assay using recombinant histone chaperones. (A) Schematic representation of the RhIP assay. The RhIP assay was performed with H3.1-H4, in the presence of either the cellular extract, or the histone chaperones Nap1 or Asf1. (B) Purified human Nap1 and Asf1 were analyzed by SDS-16% PAGE with Coomassie Brilliant Blue staining. Lane 1 indicates the molecular mass markers, and lanes 2 and 3 indicate Nap1 and Asf1, respectively. (C) RhIP-immunostaining images of H3.1. The exogenously added H3.1-3HA-H4 complex was stained with an anti-HA antibody. Cells in S phase were monitored with Cy5-dUTP. Bar indicates 10 μm.

**Figure 2- figure supplement 1.**
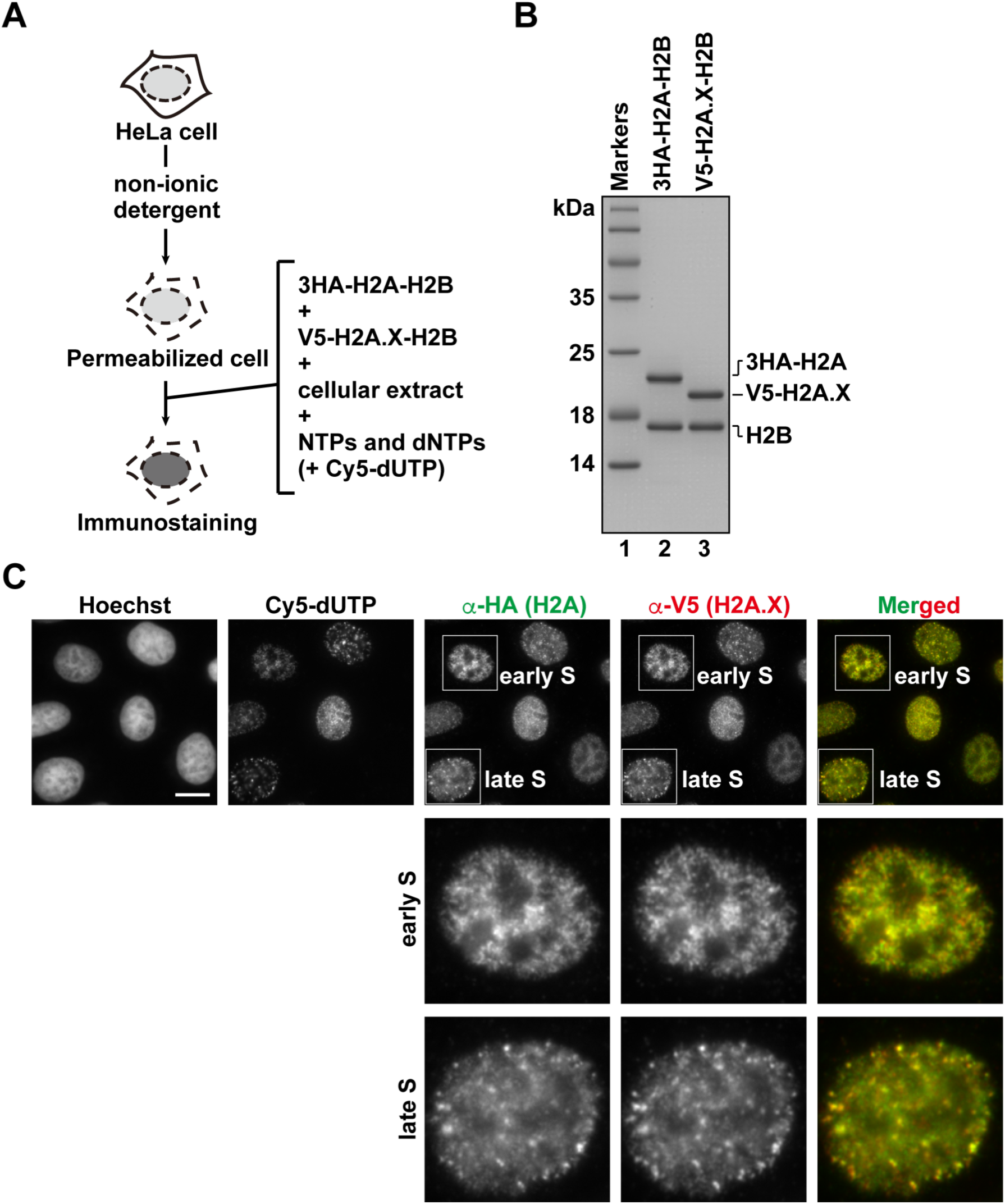
Analysis of histone H2A.X incorporation by the RhIP assay. (A) Schematic representation of the RhIP assay, using the reconstituted H2A-H2B and H2A.X-H2B complexes. (B) Reconstituted H2A-H2B and H2A.X-H2B complexes were analyzed by SDS-16% PAGE with Coomassie Brilliant Blue staining. The 3HA and V5 tags were fused to the N-termini of H2A and H2A.X, respectively. Lane 1 indicates the molecular mass markers, and lanes 2 and 3 indicate the H2A-H2B and H2A.X-H2B complexes, respectively. (C) RhIP-immunostaining images of H2A and H2A.X. Exogenously added H2A-H2B complexes were stained with an anti-HA or -V5 antibody. Cells in S phase were monitored with Cy5-dUTP. Bar indicates 10 μm.

**Figure 3-figure supplement 1.**
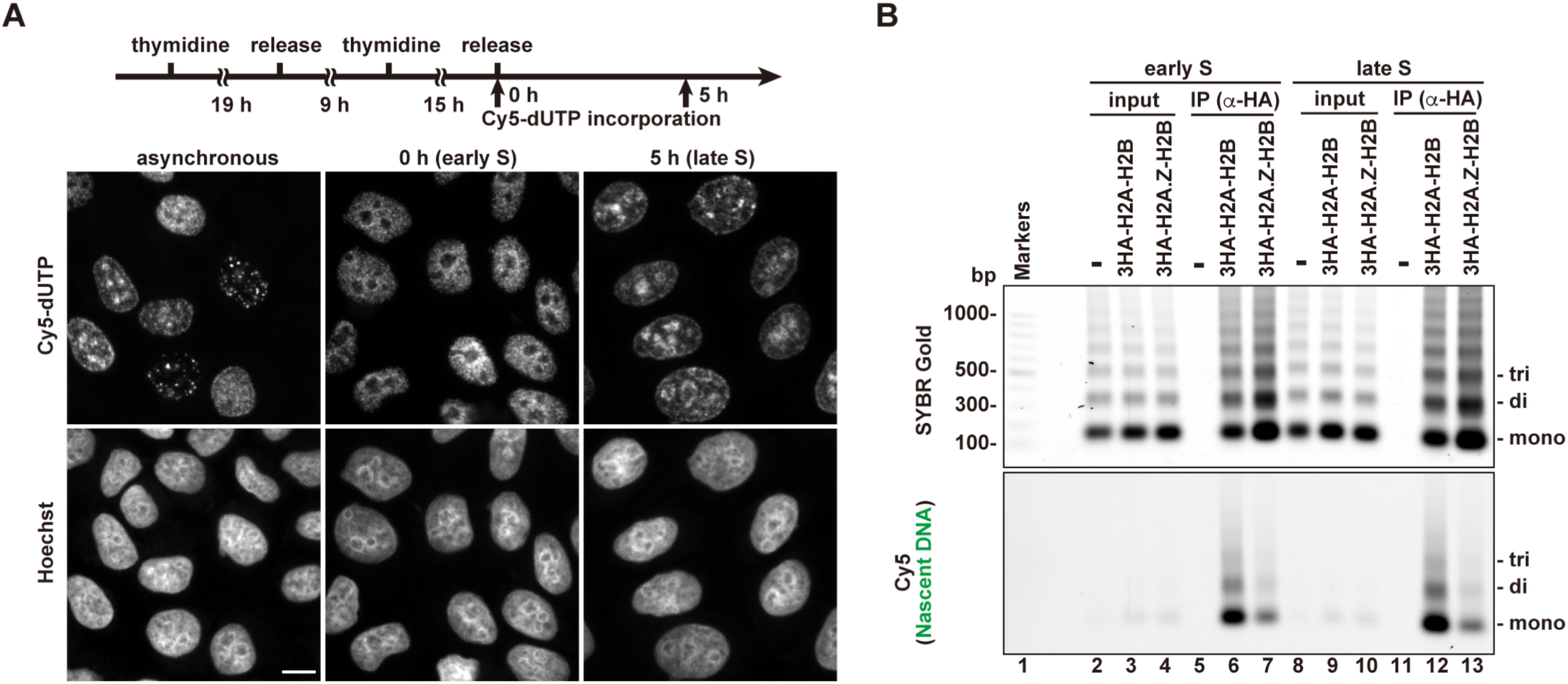
Analysis of histone H2A and H2A.Z incorporations in S phase cells with the RhIP-ChIP assay, using synchronized cells. (A) Cells were synchronized in early or late S phase by a double thymidine block (upper). Cell synchronization was confirmed by the homogeneous replication pattern, using Cy5-dUTP labeling (top images). (B) The immunoprecipitated DNA was analyzed by 1.5% agarose electrophoresis. Upper and lower images were obtained from the same gel. The DNA was visualized with SYBR Gold (upper) and the nascent DNA was visualized by detecting the Cy5 signals (lower). Lane 1 indicates a 100 bp DNA ladder. Lanes 2-7 and 8-13 indicate early S and late S phase samples, respectively. Lanes 2-4 and 8-10 indicate input samples, and lanes 5-7 and 11-13 indicate immunoprecipitated samples, respectively. Each lane indicates a negative control without the reconstituted histone complex (lanes 2, 5, 8, and 11), the samples with 3HA-H2A-H2B (lanes 3, 6, 9, and 12), and the samples with 3HA-H2A.Z-H2B (lanes 4, 7, 10, and 13).

**Figure 4-figure supplement 1.**
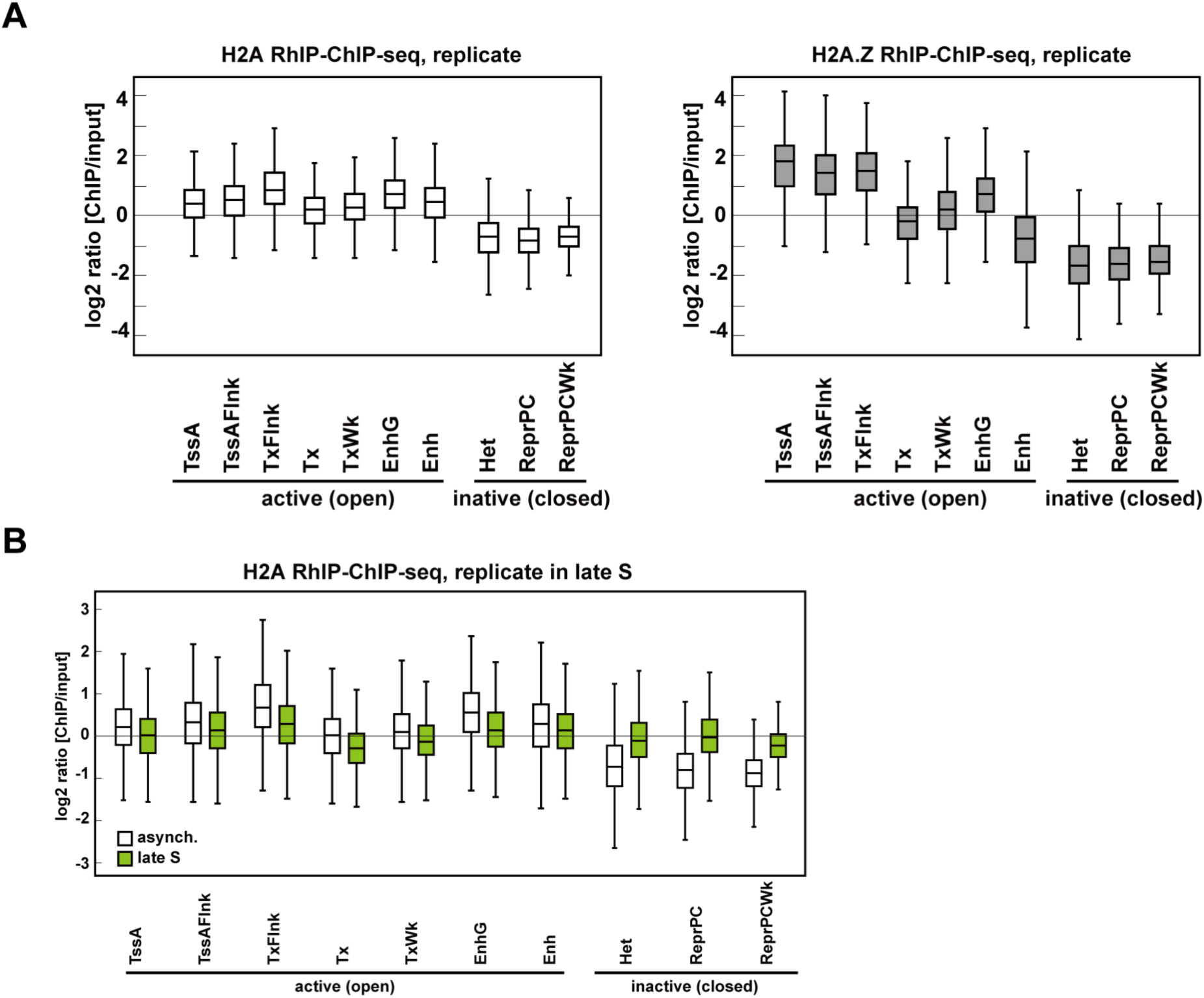
RhIP-ChIP-seq analysis of H2A and H2A.Z. (A) Biological replicate of Figure 4C. (B) Biological replicate of Figure 4E.

**Figure 4-figure supplement 2.**
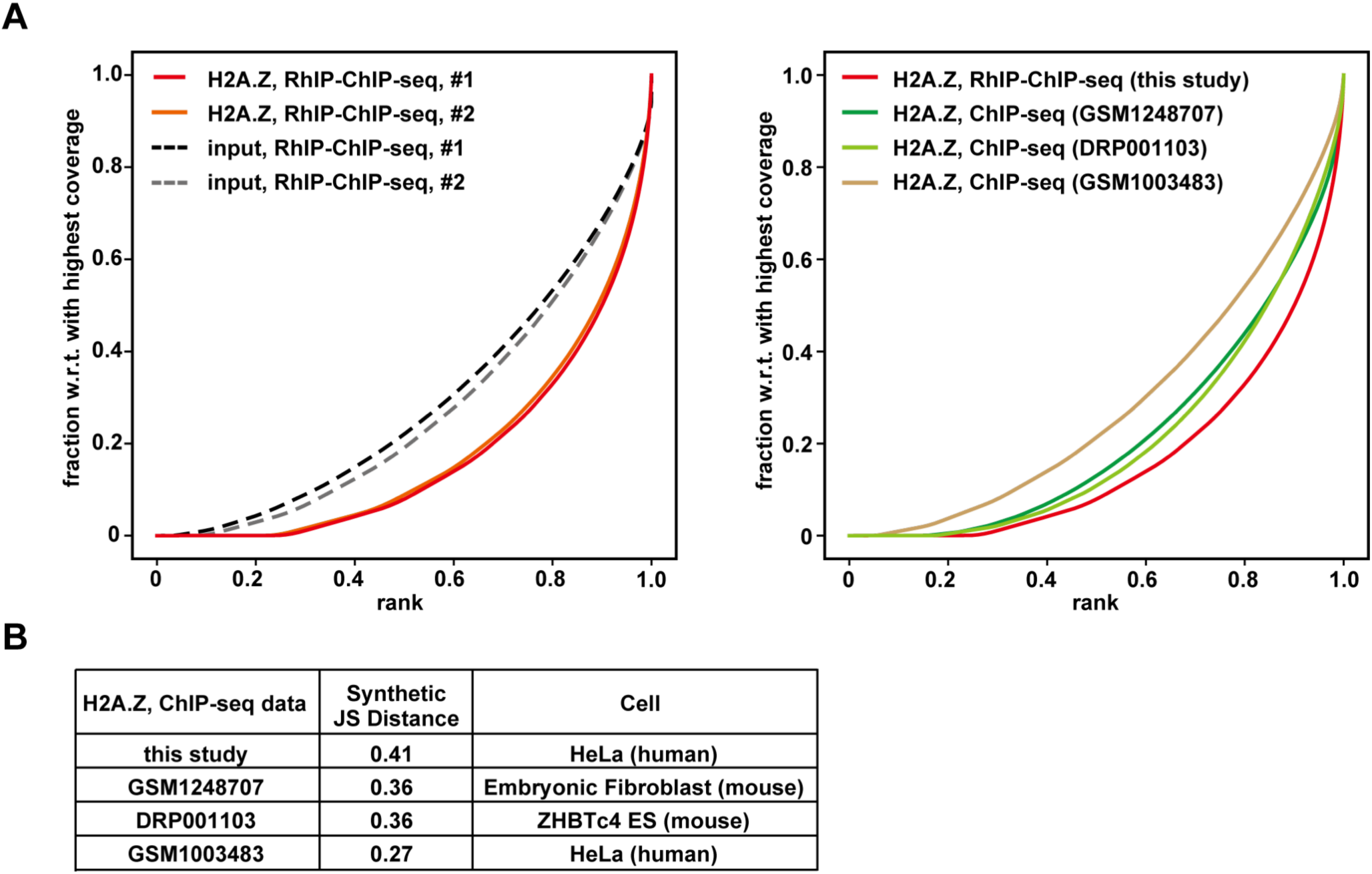
Quality check of RhIP-ChIP-seq analysis of H2A.Z. (A) The plotFingerprint of RhIP-ChIP-seq (left and right) or ChIP-seq of H2A.Z (right). (B) Quality control metrics of each sequencing data set shown in (A). The synthetic JS distance is the Jensen-Shannon distance between a given sample and the expected distribution of a perfect input sample. Higher values indicate greater difference between the two curves, with minimum and maximum values of 0 and 1, respectively.

